# Integrating ecological genomics and eco-evo-devo reveals multiple adaptive peaks in ant populations of the Arizona Sky Islands

**DOI:** 10.1101/045419

**Authors:** Marie-Julie Favé, Ehab Abouheif

## Abstract

Uncovering the genetic basis of adaptation is a great challenge facing evolutionary biologists. We ask where is the locus of adaptation from the perspective of ecological genomics (ecogen) and evolutionary developmental biology (evodevo). Ecogen focuses on identifying loci under selection between populations in different environments by scanning genome-wide patterns of genetic divergence, while evodevo focuses on candidate developmental regulatory genes and networks underlying phenotypic differences between species and higher taxa. We attempt to reconcile these different perspectives by studying the response of ant populations to past climate change on the Arizona Sky Islands - high elevation mountain ranges that represent a replicated natural experiment. We previously used an evodevo approach to show that adaptation to climatic changes in the Arizona Sky Islands in the ant species *Monomorium emersoni* occurred through repeated changes within the gene network underlying the development of alternative dispersal phenotypes: winged and wingless queens. Here, using an ecogen approach we uncovered several loci under positive selection that associate with habitat temperature. These temperature-associated loci show a repeated increase in frequency following climatic changes on each of the Sky Islands. Surprisingly, gene flow between locations within a Sky Island is restricted by temperature adaptation along the ecological gradient and not by dispersal phenotype. This finding suggests that determination of winged and wingless queens may be developmentally plastic, and this plasticity may facilitate jumps between adaptive peaks on a fitness landscape. Integration of evodevo and ecogen reveals multiple adaptive peaks and predictability at multiple biological levels within a single species.

## Introduction

Evolutionary adaptation is key for species persistence in the face of environmental changes and discovering its genetic basis has become a central focal point of modern evolutionary biology (*1*–*5*). Different approaches have been taken by its practitioners to identify the genes underlying the evolution and persistence of adaptive phenotypes across different environments (*6*–*13*). In recent years, two parallel yet opposing approaches have emerged from different sub fields: ecological genomics (*14*–*16*) and eco-evo-devo (*6*,*17*,*18*). Both of these subfields aim at discovering genes that underlie phenotypic adaptation to understand the interplay between an organism’s genes, its phenotypes, and the ecological selective forces at play. Ecological genomics takes a bottom-up approach, where regions of the genome affected by ecological selection pressures are first identified by finding genomic regions showing patterns consistent with divergent natural selection across environments acting on the genome (*14*, *16*). These regions can subsequently inform about ecologically important factors the population is adapting to. On the other hand, eco-evo-devo takes a top-down approach by first focusing on adaptive phenotypes expressed differently across environments, and then investigating candidate genes and gene networks that are likely to be involved in the generation of the adaptive phenotypic traits (*6*, *19*, *20*).

Both ecological genomics and eco-evo-devo have their own inherent strengths, weaknesses and biases, and the identification of the locus of adaptation by either of these approaches may lead to different outcomes. An ecological genomics approach allows for the discovery of natural selection signals in the genome without a priori expectations of the trait showing divergence between environments (*16*, *21*, *22*). For example, genomic regions have been identified for color polymorphism in mouse coat color (*23*, *24*), for reduction in armor plates number in sticklebacks (*13*) and for wing color patterns in Helioconius butterflies (*9*, *25*). By comparing populations of the same species living in divergent ecological conditions, genomic islands of divergence can be identified using markers distributed along the genome, such as restriction-site based methods (RFLPs, AFLPs, rad-seq), microsatellites, or single nucleotide polymorphisms (SNPs). The frequencies of the markers present in these islands of divergence can then be associated with ecological gradients from where the species have been sampled (*26*–*28*). If the precise location in the genome of the markers are known, candidate genes potentially involved in the adaptation can be identified and their function experimentally tested in the lab (*29*–*31*). The ecological genomic approach is very powerful to identify the genomic basis for traits (*1*) that have experienced a relatively hard selective sweep (*22*, *32*); (2) for which the effect of recombination is smaller than that of selection (*22*, *33*); and (3) that have a genomic architecture that allows for the identification of the genes under selection - for example, those with a strong genetic influence, and a reduced environmental contribution. Ecological genomics has been largely successful to identify the genetic basis of adaptive traits with a strong genetic basis, and for which a limited number of genes contribute to the trait variation. However, this approach faces similar challenges to genome-wide association studies (GWAS), where the genetic basis of traits with a large contribution of the environment (plastic traits), traits showing gene-by-environment interactions, and traits for which a large number of genes of small effect contribute to the phenotypic variation is difficult without investigating extremely large cohorts of individuals (*34*–*36*).

On the other hand, eco-evo-devo primarily aims at investigating phenotypes known to be adaptive because of their striking divergence between environments (*17*, *37*, *38*). For example, evolution of genes involved in adaptive changes between closely related species have been identified for leg length in semi-aquatic bugs (*7*, *39*), eye-spots in butterflies (*6*, *20*) and flower color (*8*). Leveraging the large knowledge from related model organisms, one can identify candidate genes which are known to be involved in the development of the trait from homology of gene sequences and functions. Furthermore, candidate genes can also be identified through comparative transcriptomics, together with established functions of those genes in model species (*40*–*42*). It is then possible to reveal in a favorite organism gene expression levels, spatial expression patterns, and eventually perform functional tests to determine whether and how these candidate genes have been modified in their expression and function to give rise to novel adaptive phenotypes. This approach is highly powerful to uncover genetic mechanisms underlying phenotypic adaptation and focuses immediately on genes that are likely to be involved in the development of the trait. Moreover, it gives the possibility to study not only the genetic basis of the trait, but also its dynamic changes during development, revealing changes that may not be readily observable at the adult stage (*42*, *43*). Eco-evo-devo has the power to study traits for which the environment plays an overwhelming role in the phenotypic variation, such as horns in horned beetles (*18*) or castes in hymenopteran insects (*43*–*45*). However, eco-evo-devo often limits itself to the study of (*1*) large morphological consequences often resulting from the action of a few genes of large effect, (*2*) traits and species that can be experimentally manipulated, and (*3*) relies on the assumption of the conservation of function between homologous gene sequences between related species, which may or may not be true depending on the trait and organism (*46*). Eco-evo-devo may therefore be less powerful to detect adaptive physiological traits and traits controlled by many genes of small effect.

Both approaches have been successful in revealing the genetic basis of adaptation in the wild (*47*, *48*), but unfortunately very few systems have been studied using both approaches simultaneously. Therefore, more studies integrating both approaches are needed and only by combining both, we may reveal a more complete picture of the locus of adaptation. Here we ask where is the locus of adaptation from the perspective of ecological genomics and evolutionary developmental biology using a system of replicated populations of the ant *Monomorium emersoni* inhabiting five Arizona Sky Islands (*49*).

### The Arizona Sky Islands

The Arizona Sky Islands (Fig.1A and B) are high elevation mountain ranges isolated from each other by large areas of desert (*50*, *51*). Each Sky Island is a replicate of an environmental gradient encompassing semi-desertic habitat in the low elevations, temperate forest at the mid-elevation, and coniferous forests at the highest elevations. These heterogeneous environments provide tremendous opportunities for diversification and speciation to occur in a wide variety of organisms. Its key location between the Neotropical and Neartic biomes resulted in biodiversity not exceeded anywhere else in terrestrial North America, which is reflected in the presence of numerous endemic species (*51*). Several ecological factors vary rapidly along these replicated elevational and ecological gradients, in particular, temperature, precipitation, solar radiation, habitat productivity, and habitat fragmentation. Environmental changes that would normally be observed along thousands of kilometers of latitudinal gradients are observed along only a few hundred meters of elevation change. This imposes unique adaptive challenges for organisms whose distributional ranges encompass more than one elevation zone. Indeed, dispersal of an organism over only a few kilometers may impose completely different ecological conditions and thus challenge reproduction and survival.

**Figure 1:**
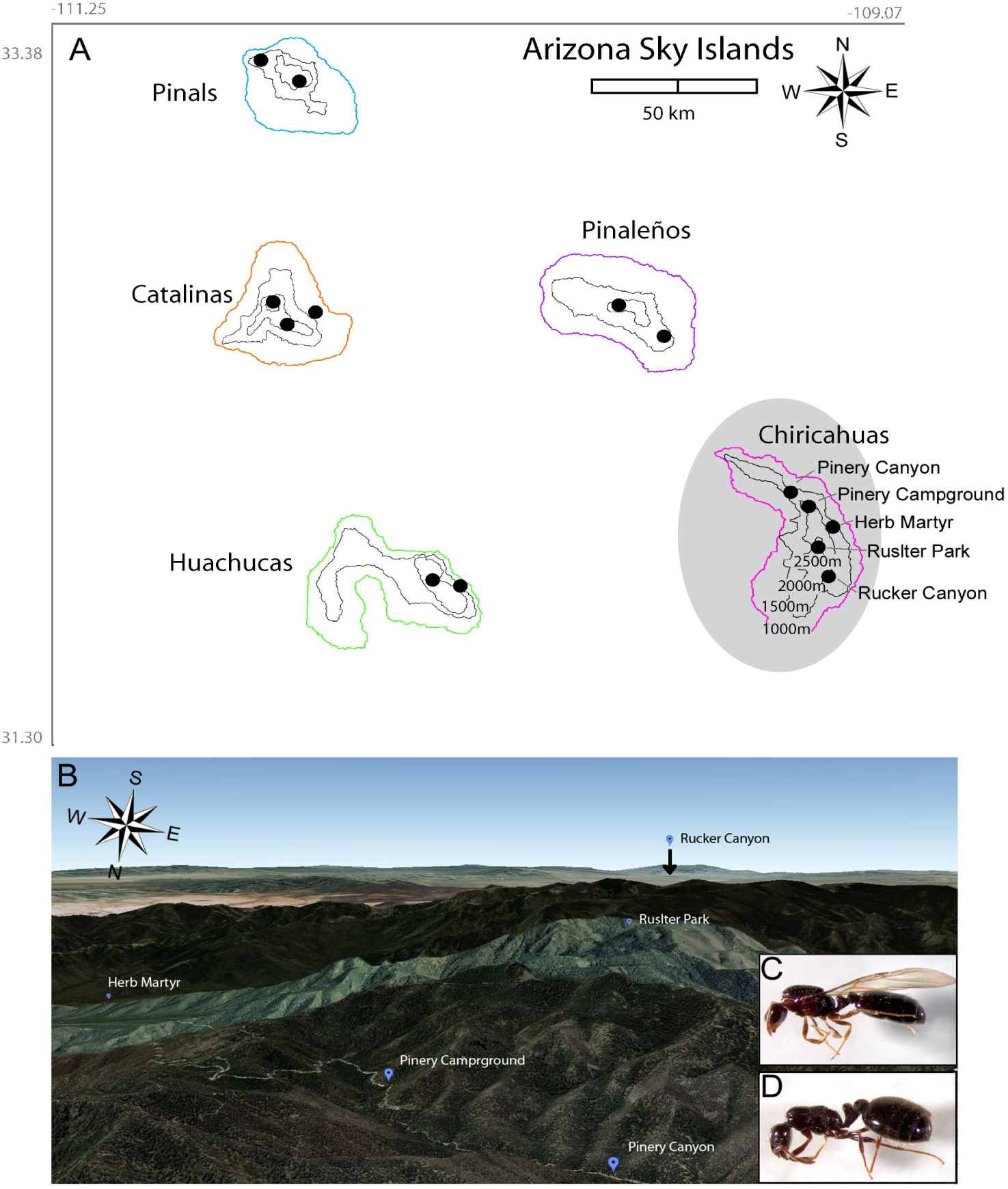
M. *emersoni* populations in the Arizona Sky islands. (A) Map of the Arizona Sky Islands, modified from Fave et al (2015) (*49*)(B) Aerial map of the Chiricahuas Sky Island and sampling sites. *Monomorium emersoni* queens, (C) winged and (D) wingless.

### Parallel loss of wings in queens of *M. emersoni*

Two *M. emersoni* queen phenotypes co-exist: winged and wingless (Fig.1C and D). While the diversification of the queen caste across ants has been recognized for a long time (*52*, *53*), its functional and evolutionary importance has only been recently embraced (*54*–*56*). Alternative phenotypes in ant queens differ not only in their dispersal abilities and morphologies, but also in their resource allocation during colony founding. During this key life history stage, winged queens hydrolyze their flight muscles for energy, whereas wingless queens rely on workers that disperse along with them from the mother colony to provide help and resources. This life history change has evolved multiple times independently in all major ant sub-families and is therefore considered to be an adaptive trait (*54*–*56*).

*M. emersoni* colonized each Sky Island independently from an ancestral continuous population, but nowadays, populations are isolated on each Sky Island without gene flow occurring between them (*49*). Inference of demographic changes in *M. emersoni* populations indicated the existence of a continuous population around 100,000 years ago, which separated into a Northern and a Southern groups around 80,000. Eventually each Sky Island population got isolated from one another following the warm up after the last glacial maximum (*49*) (Summarized in Fig. 2). In *M. emersoni*, the wingless phenotype evolved in parallel, and occurs at higher frequencies at high elevations and in fragmented habitats on each of the Arizona Sky Islands, suggesting that wingless queens repeatedly and independently adapted to these elevation gradients on each of the Sky Islands (*49*).

**Figure 2:**
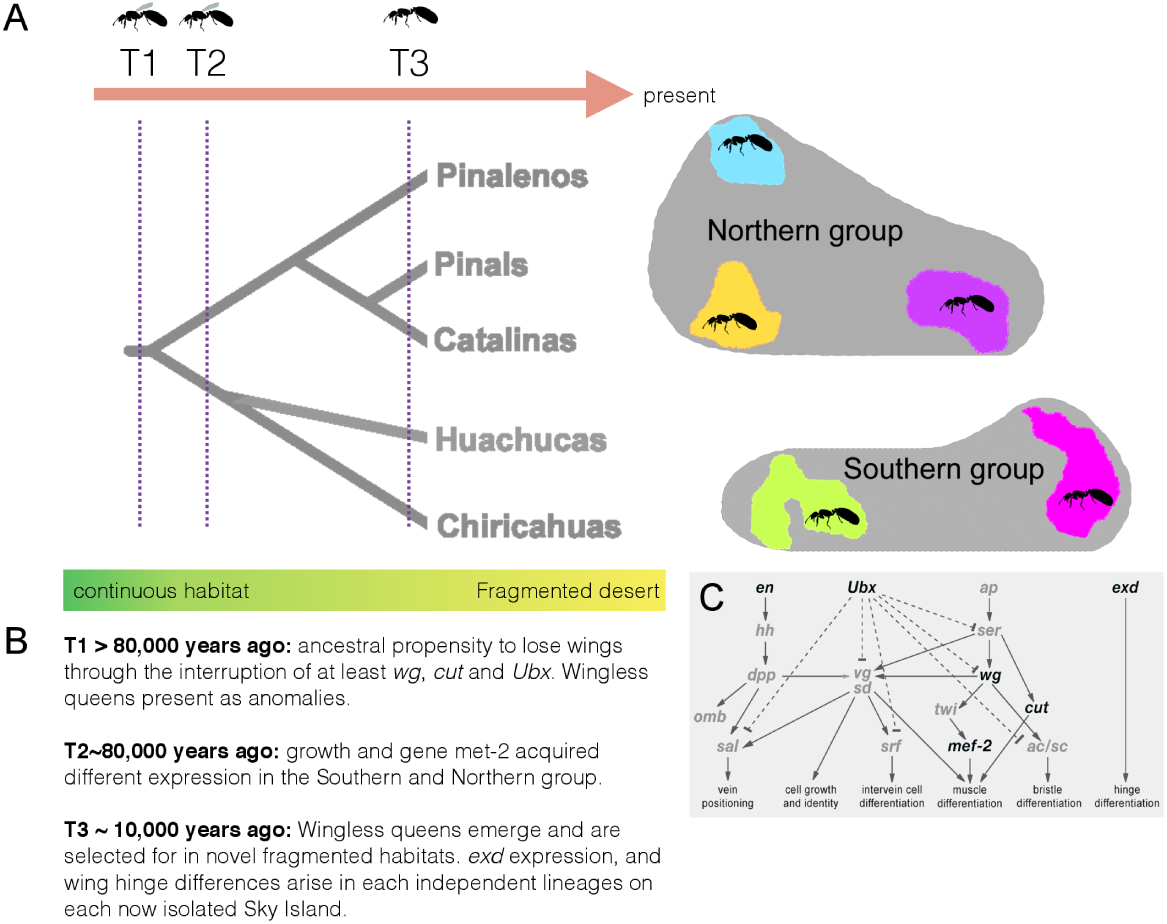
Summary of climatic changes, population history, wingless phenotype evolution, and its associated changes in gene regulatory network and other phenotypes. (A) The population tree represents the demographic history during the last 200,000 years. The time points T1, T2, and T3 represent key event during the evolution of the populations of *M. emersoni* and are described in (B). The geographic location of the Northern and Southern Sky Islands is depicted on the right. (C) Gene regulatory network underlying wing developement is conserved in ant queens relative to Drosophila. Bold gene names are the genes that were studied in (*49*)

Wing development in insects has become a model system to study gene interaction networks, and the deep knowledge of the wing patterning network in *Drosophila melanogaster* and other holometabolous insects has helped in identifying candidate genes involved in wing development in ants (*43*–*45*, *57*). The evolution of the wing patterning gene network between the 5 replicated populations of *M. emersoni* on Sky Islands (Fig. 1A) has been studied previously (*49*). Each population has independently evolved the loss of wings from a once unique ancestral population followed by the evolution of high frequencies of winged queens. Wings originate in queen larvae as imaginal discs, small synchronized groups of cells that will become wings during metamorphosis. Although lacking wings as adults, wingless queens still produce rudimentary or vestigial discs as larvae (*49*). In wingless queens, wing vestigial imaginal discs express a certain number of genes of the wing patterning gene network, while the expression of other genes is completely absent (*49*). The genes for which expression is completely lost in wingless queen larvae show a similar loss of expression across the five populations. Such genes, showing similar expression patterns across populations, suggest predictability in the evolution of gene networks across different replicates. On the other hand, other genes are also expressed, but their pattern of expression is different between the Northern and Southern populations (*49*). Interestingly, these groups of populations also reflect the demographic history patterns of the populations, suggesting that the expression differences between populations were established after their separation. For example, the expression of *mef2,* a key gene for wing muscle development, differs between wingless-queen larvae from Northern and Southern Sky Islands populations (Fig. 2), and shows a signature of the ancient demographic split between these two groups (*49*). Another gene *exd* for which wingless-queen larvae from each Sky Island exhibit population-specific expression patterns (*49*), not related to ancient demographic history. This suggests that these differences arose following the complete isolation of the five Sky Island populations around 10,000 years BP and are therefore an indirect consequence of climatic warming since the last ice age.

In addition to elevation and habitat fragmentation, *M. emersoni* populations have responded directly to climatic changes by evolving a temperature tolerance in warmer habitats (*49*). Heat shock and chill-coma recovery of *M. emersoni* individuals coming from different habitats showed that *M. emersoni* colonies coming from warm sites (usually lower elevations habitats) can resist a heat shock much longer than colonies originating from cooler habitats. The reverse is also true - colonies from cooler sites recover much quicker from a cooling episode than colonies coming from warmer sites. Temperature tolerance in *M. emersoni* is therefore variable between sites with different temperature regimes, and this observation suggests that *M. emersoni* temperature adaptation in the Sky Islands may be a response to the Holocene climatic warm up. Given the former cooler climate across the region, the tolerance to heat shock likely evolved repeatedly on each Sky Island as an adaptation to the warming climate at lower elevations after the last glaciation.

Therefore, using an eco-evo-devo approach, we established that the evolution of wingless queens in the Arizona Sky Islands represents an evolutionary novelty that arose following a climatic and environmental change: the gain of a developmental switch to generate the wingless queen phenotype, the complete loss of wings and wing muscles, multiple alterations in the expression of the core developmental network underlying wing development, and the gain of a new life history strategy. The repeated wing loss along each ecological gradient provides support for a predictable and deterministic outcomes of evolution when organisms encountered similar selection pressures, however, this predictability is accompanied with slight differences in gene expression patterns between populations that reflect historical demographic events. This eco-evo-devo approach revealed key genetic and phenotypic events that happened during the adaptation of *M. emersoni* in terms of its dispersal abilities when colonizing steep environmental gradients. However, dispersal abilities may not be the only adaptation that allowed *M. emersoni* cope with the changing conditions.

Here, in an effort to combine eco-evo-devo with ecological genomics, we use a bottom-up approach to uncover whether and how natural selection has shaped *M. emersoni* genome. Ecological selective forces during colonization of novel habitats are numerous, and may produce complex patterns of adaptations on both the genotype and the phenotype. By using a combination of eco-evo-devo and ecological genomics approaches, we reveal that *M. emersoni* populations show genomic signatures consistent with temperature adaptation, that this recurrent adaptation is shaping gene flow patterns within Sky Islands. Our data raise the possibility that the evolution and maintenance of the dispersal phenotype of *M. emersoni* queens and its associated gene network may have been facilitated by phenotypic plasticity.

## Material and Methods

### Large scale sampling

We collected *M. emersoni* colonies from 14 sites distributed along the elevation gradients of five Sky Island mountains in Southeastern Arizona, USA (Fig. 1, Table 1). For each site, we measured the latitude and elevation in the field, and we obtained the average annual temperature, temperature seasonality, annual precipitation from WorldClim Atlas and the vegetational growth average were obtained from the National Atlas (Table 1). Both databases have a 1 km2 resolution. Our personal observations confirmed the databases are of the same magnitude as the area of our average sampling sites. Ecological gradients along each Sky Island do show similar trends, but depending on their altitude and their latitudinal location, also show variation in their absolute values. For example, our sites from the Huachucas show, on average, higher annual temperatures (Table 1).

**Table 1:**
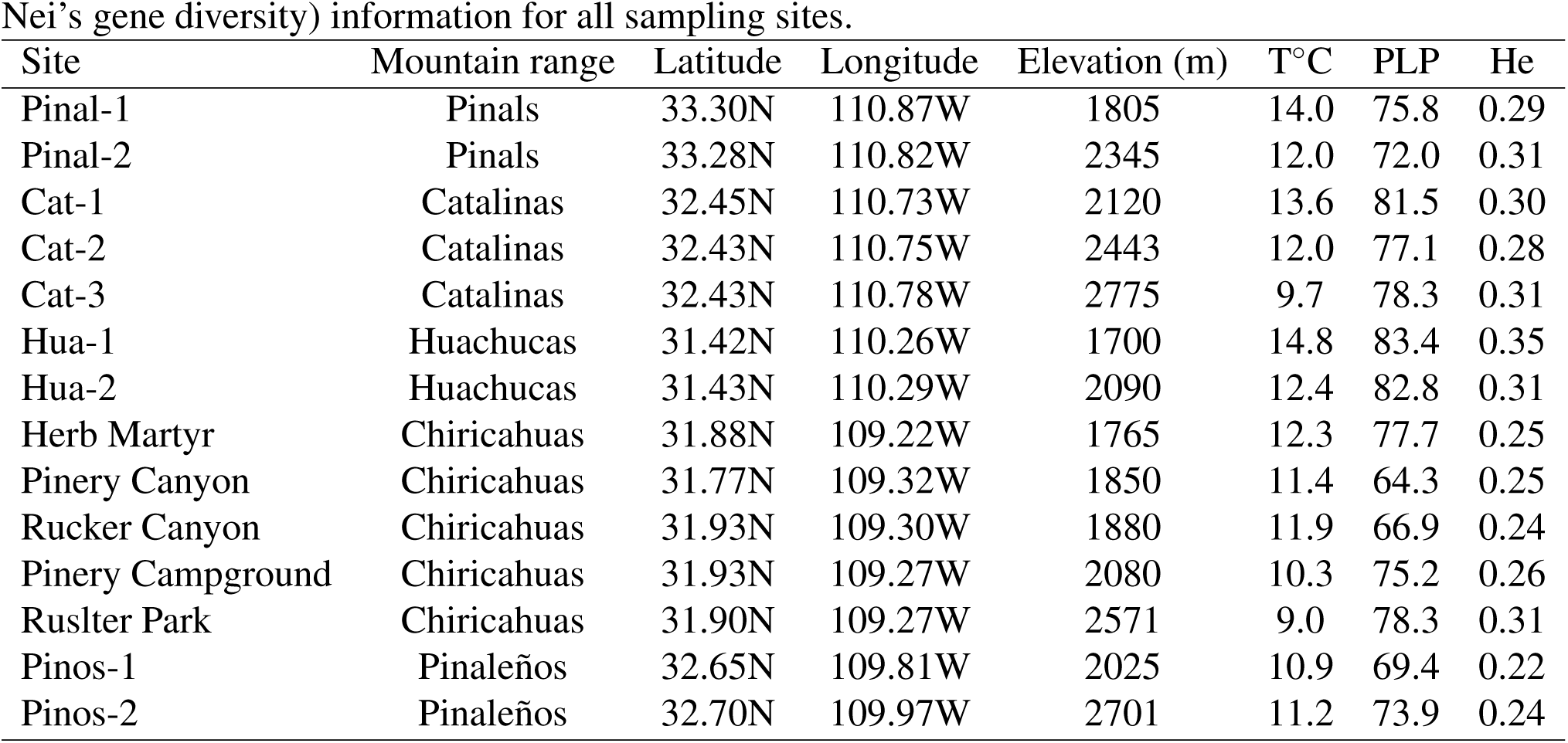
Locations of sampling sites. Fourteen sites have been sampled across the Arizona Sky Islands. Geographic (Latitude, Longitude, Elevation), environmental (T°C: temperature) and genetic (PLP: percentage of polymorphic loci, He: Nei’s gene diversity) information for all sampling sites.

### Small scale sampling

For our ecological genomics approach, we selected the Chiricahuas Sky Island (Fig. 1) as our focal Sky Island, and performed a deeper sampling of the ecological gradient and landscape. We collected colonies from five distinct sites that represent three different types of habitats: Herb Martyr is situated in lower mid-elevation area, dominated by junipers and subtropical species such as agave and yucca. Pinery Canyon and Rucker Canyon are situated on opposite sides of the mountain, but are similar in their habitat, dominated by oak-juniper woodlands. Pinery Campground is geographically close to Pinery Canyon, at a higher elevation, but is dominated by Douglas pines and species usually found at higher latitudes. The highest site, Rustler park, is also dominated by subalpine species such as Douglas fir and Pinyon pines. Our sampling design in the Chiricahuas consists therefore of an altitudinal and ecological gradient both spanning a defined geographic area (Fig. 1). We specifically included sites similar in their habitat but geographically far apart on the mountain as well as sites situated in close vicinity but different in their habitat. This design allows for the discrimination between geography or habitat as being the dominant factor in shaping neutral and adaptive divergence in *M. emersoni* and investigate the effects of the ecological gradients on gene flow patterns.

### DNA extraction and genotyping

We chose only one individual queen per colony for genetic analyses to avoid pseudo-replication of the genetic data as ants from the same colony are generally closely related individuals. We extracted DNA from 318 *M. emersoni* queens, using thoracic tissue from preserved individuals. We genotyped all individuals for Amplified Fragment Length Polymorphisms (AFLP) markers, as described in (*58*). We used four primer combinations to generate 157 unambiguous loci: MCTA and EACC, MCTA and EACG, MCAG and EACC, MCAG and EACG. We ran the PCR products on a LiCor DNA analyser 4300 and the same researcher scored them blindly using the accompanying Saga software. About 10% of the samples were amplified and scored twice to estimate the error rate in the procedure, which is similar to what was reported in other studies (*59*).

### Identifying natural selection in *M. emersoni*

To identify regions of the genome of *M. emersoni* that are under natural selection, we identified outlier loci using three different methods, each of which is based on a different framework. We first used the method of (*60*) modified for dominant markers (dFdist) and implemented in Mcheza by (*61*). This approach is based on the assumption that loci under divergent selection will display significantly higher Fst values than the majority of loci that are evolving neutrally or near-neutrally. Loci under stabilizing selection will display lower than expected Fst values. This model compares the empirical distribution of Fst values to a simulated distribution of Fst values conditional on heterozygosity in a subdivided population under the infinite island model. We performed global comparisons over all 5 Sky Islands, pairwise comparisons between Sky Islands, as well as global and pairwise comparisons between the 5 Chiricahuas sites. We did not retain any loci with lower than expected Fst because divergence-based methods have little power to detect stabilizing selection (*60*, *62*). Second, we used the method of (*63*) implemented in Bayescan to detect outlier loci. Bayescan uses a Bayesian method to directly estimate the probability of each locus to be under divergent selection. It is considered to be more robust to deal with complex demographic scenarios, but is also considered to be more conservative in outlier identification (*63*). We therefore used the Bayescan approach using data from the 5 Sky Islands to confirm our outlier identification from dFdist. We used a third approach to gain insights into the genomic adaptive divergence in *M. emersoni* by identifying loci that co-vary with environmental parameters. We used the approach of (*64*) implemented in SAM which performs regressions between allele frequencies and environmental variables to identify candidate loci potentially under natural selection. We tested for association with annual average temperature, temperature seasonality, annual average precipitation level, and habitat vegetation growth. This approach is complementary to Fst or Bayesian outlier detection and associates polymorphisms directly with environmental factors. Therefore, if loci previously identified with Fst or Bayesian methods also show an association with environmental factors, the putative adaptive nature of these loci is further supported, and a particular ecological pressure can be suggested as being responsible for the signals of natural selection.

### Linkage disequilibrium

We inferred the genomic distribution of outlier loci across the genome by testing for the presence of linkage disequilibrium among them. The presence of a higher linkage disequilibrium among outlier loci than among randomly selected neutral loci would indicate that outlier loci are likely to be physically linked to each other in the genome, or that there is population substructure. We tested for the presence of linkage disequilibrium among the outlier loci using the program DIS (*65*), using the maximum-likelihood based method equivalent to that of (*66*).

### Population diversity and structure

After the identification of outlier loci, we removed these loci from the data set and retained only putatively neutral markers for following analyses: allele frequencies were estimated with AFLP-SURV (*67*) using the Bayesian approach of (*68*) for dominant markers. We calculated the proportion of polymorphic loci, expected heterozygosity and average gene diversity for all populations. We calculated pairwise Fst estimates between sites using all loci, and on the putatively neutral loci data set only.

Within the Chiricahuas, we also reveal genetic structure among sites with STRUCTURE (*69*, *70*). Using multilocus genotypic data, STRUCTURE attempts to form groups of individuals without any a priori information on their origin. We performed the admixture algorithm for 100 000 iterations, plus a burn-in period of 100 000 iterations. We determined the most likely number of clusters, K, as the number reached when increasing K further did not show any improvement in population structuring (*71*). We then estimated for each K cluster identify by STRUCTURE the proportion of individuals coming from each site.

### Relationships between queen phenotype, habitat type, and gene flow

Queen phenotype frequencies (winged / wingless) at each site were obtained from previous work (*49*). Average queen phenotype frequencies was estimated for all pairwise comparisons within the Chiricahuas as a proxy for dispersal potential between two sites. We tested several scenarios that could be underlying the gene flow patterns within the Chiricahuas: isolation-by-distance (3D geographic distance), isolation-by-temperature (average annual temperature), and isolation-by-phenotype (winged queens). We performed Mantel tests (*72*), using 1000 replications for measuring the significance levels, with the R package vegan. We used partial Mantel tests to test whether the presence of covariates modulates the associations.

## Results

### Detection of Fst outliers

To identify regions of the genome of *M. emersoni* that show variation patterns consistent with natural selection, we performed a series of comparisons between *M. emersoni* Sky Island populations (see Methods and Table 3). Outliers were detected in all 22 comparisons at the 95% level: the Global Sky Island dataset, 10 Sky Island pairwise comparisons, global Chiricahuas dataset, and 10 pairwise comparisons between the 5 Chiricahuas sites. Using dFdist, we identified 7 outliers in the global dataset, and 13 in the pairwise comparisons between Sky Islands. The 7 global outliers were also detected in more than one Sky Island pairwise comparison (Table 3). We then replicated our analyses using only the Chiricahuas data set, in which we also perform both global and pairwise comparisons between sites. We identified 16 repeated outliers in the Chiricahuas, and among these, 7 are also replicated in the global Sky Island dataset. Therefore, based on dFdist analyses, we detected 7 repeated outliers (M1, M36, M37, M63, M115, M135 and M148) that show patterns of divergence coherent with divergent natural selection in the Arizona Sky Islands (Table 3). We then used the method implemented in Bayescan to identify outliers in the global dataset to confirm our outlier identification with dFdist. We confidently identified one outlier loci in the global dataset, which was also repeatedly identified by dFdist at both the global level and within the Chiricahuas (M37) (Table 3). Given the conservative nature of Bayescan, this finding further supports the adaptive nature of the outlier locus M37.

**Table 3:** Comparisons performed and number of outlier loci detected in each one. Comparisons and performed with the three different approaches. The loci detected in group 1, 4, 5, 6 and 7 are Ml, M36, M37, M63, Ml 15, M135 and M148. The single loci identified with Bayescan is M37.

| Group of loci | Comparison | Number of loci |
| --- | --- | --- |
| 1 | Global outliers (dFdist) | 7 |
| 2 | Repeated outliers among pairwise Sky Island comparisons (dFdist) | 13 |
| 3 | Repeated outliers pairwise comparisons within the Chiricahuas (dFdist) | 16 |
| 4 | Repeated outliers detected in both Global and within Chiricahuas comparisons (dFdist) | 7 |
| 5 | Loci associated with temperature (SAM) | 7 |
| 6 | Group 5 and associated with temperature | 7 |
| 7 | Group 6 and in comparisons involving sites with different temperature | 7 |
| 8 | Group 6 and Bayescan | 1 |

### Outlier associations with environmental factors

To identify potential environmental variables responsible for the changes in allelic frequencies in the outlier loci, we performed regressions between allele frequencies and environmental variables showing strong variation along the ecological gradients (annual average temperature, temperature seasonality, annual average precipitation level, and habitat vegetation growth). We detected 19 loci associated with annual average temperature, 13 with temperature seasonality, 4 with average annual precipitation level, and none with habitat vegetation growth. The majority of the loci associated with either temperature seasonality or precipitation levels were also associated with annual temperature. These ecological factors tend to covary with one another, and temperature is generally considered as the most important factor underlying altitudinal zonation in the Sky Islands (*73*). We found that the group of 7 repeated outliers identified with dFdist (M1, M36, M37, M63,M115, M135 and M148) and the one outlier identified with Bayescan (M37) were also found to be associated with the annual average temperature Fig.3. We therefore now refer to this group of loci as ‘temperature-associated loci’. This overlap in loci identification using three analyses build on different premises and assumptions is compelling, and supports further the possible adaptive value of these loci. Furthermore, we previously demonstrated experimentally a strong effect of temperature on *M. emersoni* survival, where colonies originating from cold sites were much more sensitive to heat shock than colonies coming from warm sites (*49*). Collectively, these results support the hypothesis that *M. emersoni* has adapted to the temperature gradient that was established during the climatic change following the Pleistocene glaciation.

**Figure 3:**
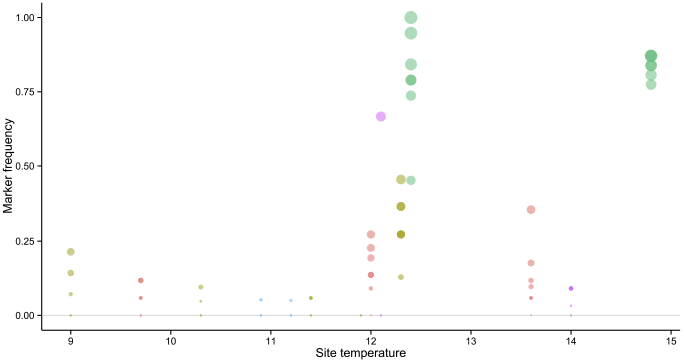
Relationship between temperature and alleles frequencies of outliers, and genetic differentiation between sites. Purple: Pinals, Red: Catalinas, Green: Huachucas, Blue: Pinaleños and Olive: Chiricahuas. Size of the bubble represent the allelic frequency. All sites with annual temperature above 12C have alleles frequencies above zero in most outlier loci, and their frequencies are generally much higher than sites with annual temperature lower than 12C.

### Temperature-associated loci identify a single region in the genome under selection

Loci identified through genome scans can either be segregating independently form each other, or physically linked to each other and tagging a single region of the genome under selection. Given a sufficient number of loci distributed randomly across the genome, association signals are often detected for several loci in physical proximity of each other and therefore are not segregating independently. We quantified the levels of linkage disequilibrium among our 7 temperature-associated loci by testing whether they segregated independently from each other. As a negative control, we also tested linkage disequilibrium in a set of 20 randomly selected neutral loci from our dataset (Fig. 4). We found that linkage disequilibrium is significantly higher among our 7 temperature-associated loci than among random neutral loci (p < 2.2 x 10e-16). This indicates that these loci segregate together and most likely tag a single region in the genome where a strong signal of selection exist (one single island of divergence). Our power to detect several islands of lower divergence is limited by our total number of loci, and therefore, other regions under selection may eventually be detected with additional loci.

**Figure 4:**
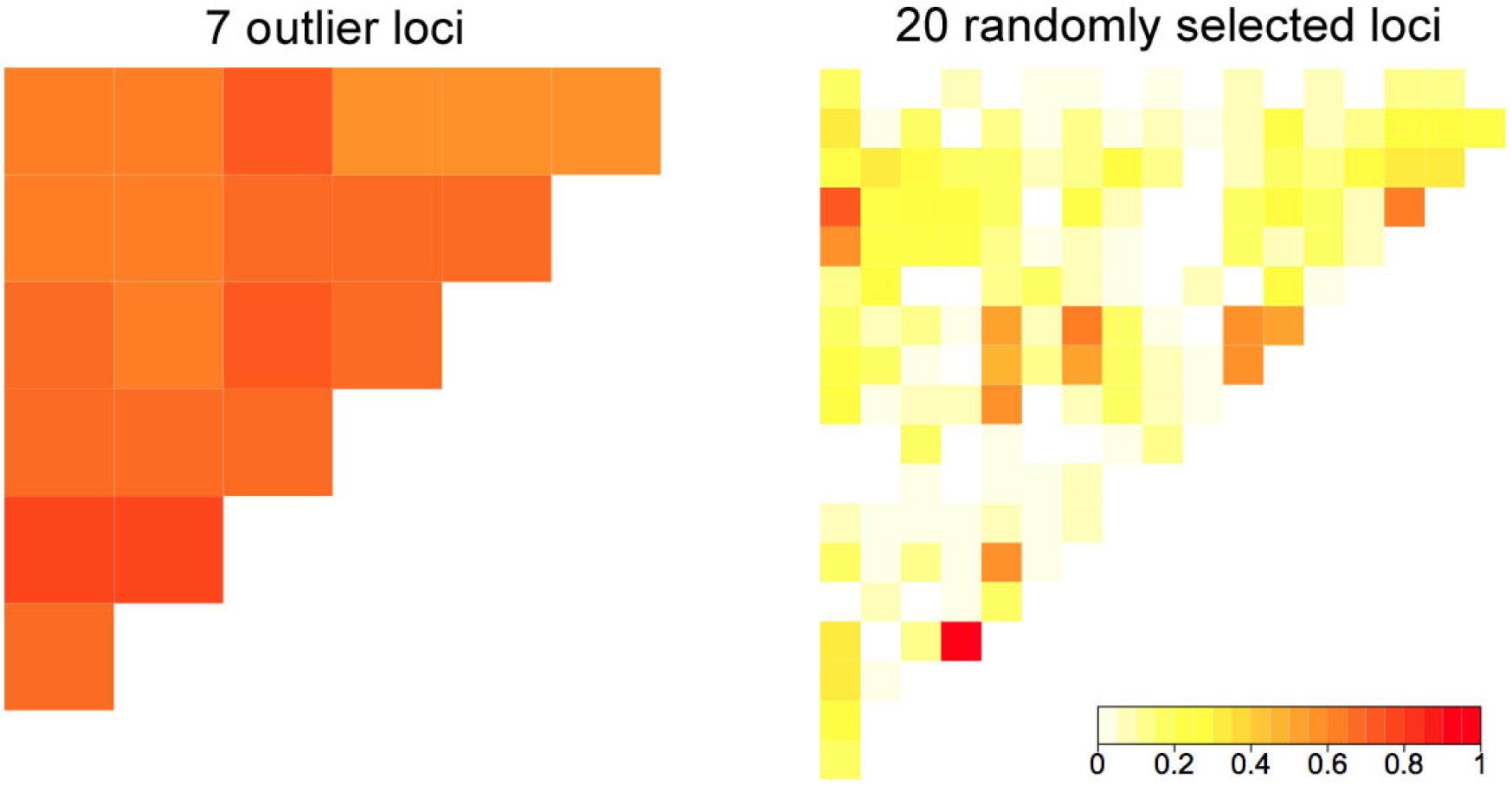
Linkage disequilibrium heat maps. Linkage disequilibrium among (A) 7 outlier loci and (B) 20 randomly neutral loci. Intensity of the color reflects the amount of linkage between two loci, white being 0, red being 1.

### Population structure and gene flow within the Chiricahuas

*M. emersoni* populations have recurrently adapted on each Sky Island to the ecological challenges imposed by the changing climate after the Pleistocene glaciation, by (*1*) evolving a novel wingless queen phenotype found predominantly in fragmented habitats at higher elevations, and by (*2*) adapting to the newly established temperature gradients of each Sky Island on both the genetic and phenotypic levels. We found that genetic divergence is much higher and significant between Sky Islands than between sites within one Sky Island (Table 2). However, Sky Islands are highly ecologically heterogeneous environments, and gene flow within a Sky Island is likely affected by trade-offs in response to ecological barriers, as often observed between ecologically divergent sites (*74*–*76*). Both adaptations we uncovered in *M. emersoni* may have affected population structure and gene flow patterns within a Sky Island. Queens must reconcile two adaptive phenotypes that respond to different ecological selective pressures: the queen wing phenotype (winged or wingless) must be adapted to fragmented and high elevation habitats, but simultaneously, the queen has to adapt to the local temperature of the habitat it disperses to. Moreover, the queen wing phenotype itself will most likely influence the dispersal of virgin queens and consequently, affect the gene flow patterns and population structure within a Sky Island.

**Table 2:**
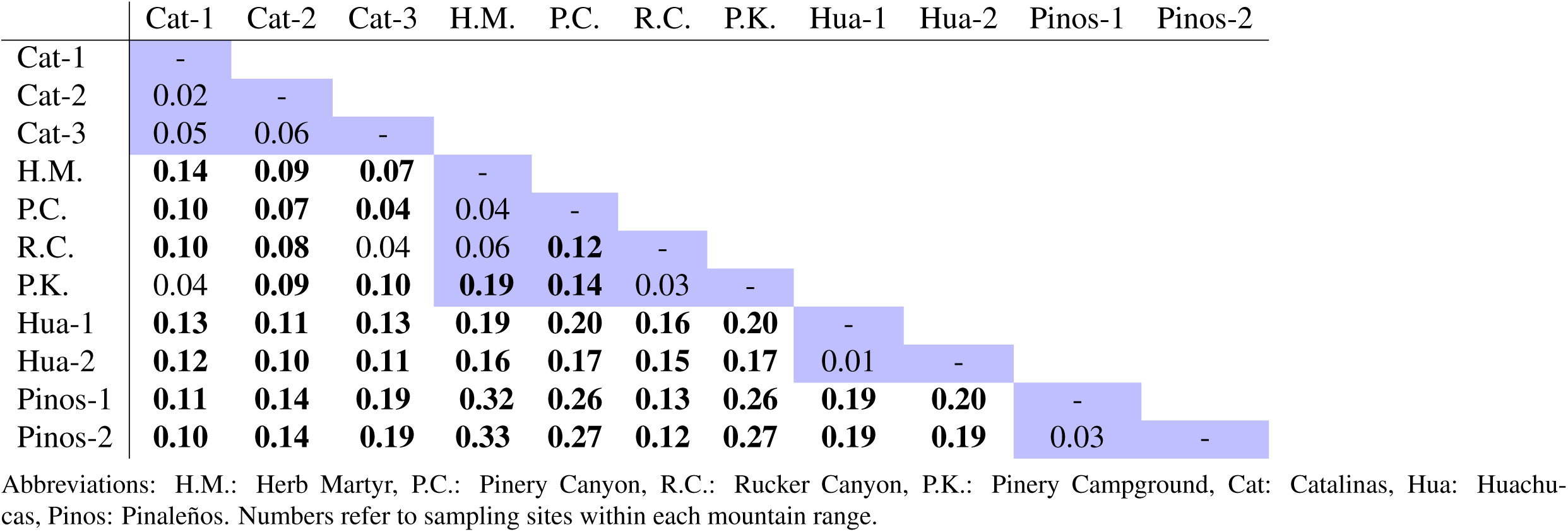
Genetic differentiation for neutral loci. Pairwise Fst estimates between collecting sites as estimated with AFLP-surv, including only putatively neutral AFLP loci. Bold values indicate a significant value after correcting for multiple comparisons (Bonferroni) and shaded values represent within-mountain comparisons.

|  | Cat-1 | Cat-2 | Cat-3 | H.M. | P.C. | R.C. | P.K. | Hua-1 | Hua-2 | Pinos-1 | Pinos-2 |
| --- | --- | --- | --- | --- | --- | --- | --- | --- | --- | --- | --- |
| Cat-1 | - |  |  |  |  |  |  |  |  |  |  |
| Cat-2 | 0.02 | - |  |  |  |  |  |  |  |  |  |
| Cat-3 | 0.05 | 0.06 | - |  |  |  |  |  |  |  |  |
| H.M. | **0.14** | **0.09** | **0.07** | - |  |  |  |  |  |  |  |
| P.C. | **0.10** | **0.07** | **0.04** | 0.04 | - |  |  |  |  |  |  |
| R.C. | **0.10** | **0.08** | 0.04 | 0.06 | **0.12** | - |  |  |  |  |  |
| P.K. | 0.04 | **0.09** | **0.10** | **0.19** | **0.14** | 0.03 | - |  |  |  |  |
| Hua-1 | **0.13** | **0.11** | **0.13** | **0.19** | **0.20** | **0.16** | **0.20** | - |  |  |  |
| Hua-2 | **0.12** | **0.10** | **0.11** | **0.16** | **0.17** | **0.15** | **0.17** | 0.01 | - |  |  |
| Pinos-1 | **0.11** | **0.14** | **0.19** | **0.32** | **0.26** | **0.13** | **0.26** | **0.19** | **0.20** | - |  |
| Pinos-2 | **0.10** | **0.14** | **0.19** | **0.33** | **0.27** | **0.12** | **0.27** | **0.19** | **0.19** | **0.03** | - |

Abbreviations: H.M.: Herb Martyr, P.C.: Pinery Canyon, R.C.: Rucker Canyon, P.K.: Pinery Campground, Cat: Catalinas, Hua: Huachuca, Pinos: Pinalenos. Numbers refer to sampling sites within each mountain range.

To reveal the population structure within the Chiricahuas, we used the STRUCTURE algorithm (*69*, *70*) on the putatively neutral loci. We found that K=3 represents the most likely number of clusters in the Chiricahuas because an increase in K did not resolve any additional clusters (*71*). To highlight similarities and differences between sites, we calculated the proportion of individuals belonging to each of the three K clusters for each of the 5 sites (Fig. 5). We found that Herb Martyr is mostly composed of individuals from K1, with certain proportion of K2 individuals. Pinery Campground is dominated by individuals from K2. Pinery Canyon and Rucker Canyon are relatively similar in their composition, with a large proportion of individuals from K3. Ruslter Park shows an admixted composition, with a relatively large proportion of individuals from K2. The similar composition of Pinery Canyon and Rucker Canyon suggests higher gene flow of individuals between them, homogenizing the genetic composition between these two sites. Furthermore, we found a correlation between the K composition of individuals at each site and the average annual temperature: Herb Martyr, the site with the highest annual temperature, shows a unique genetic composition dominated by K1 individuals, suggesting limited successful immigrant individuals into this site coming from other sites that were sampled in this study. The two sites with intermediate temperatures, Pinery Canyon and Rucker Canyon, have a similar composition dominated by individuals of K2 group, but also have some proportion of individuals form K1. The two coldest sites, Pinery Campground and Ruslter Park, both have a significant proportion of individuals with genetic background K3, suggesting a limited dispersal from other sites sampled.

**Figure 5:**
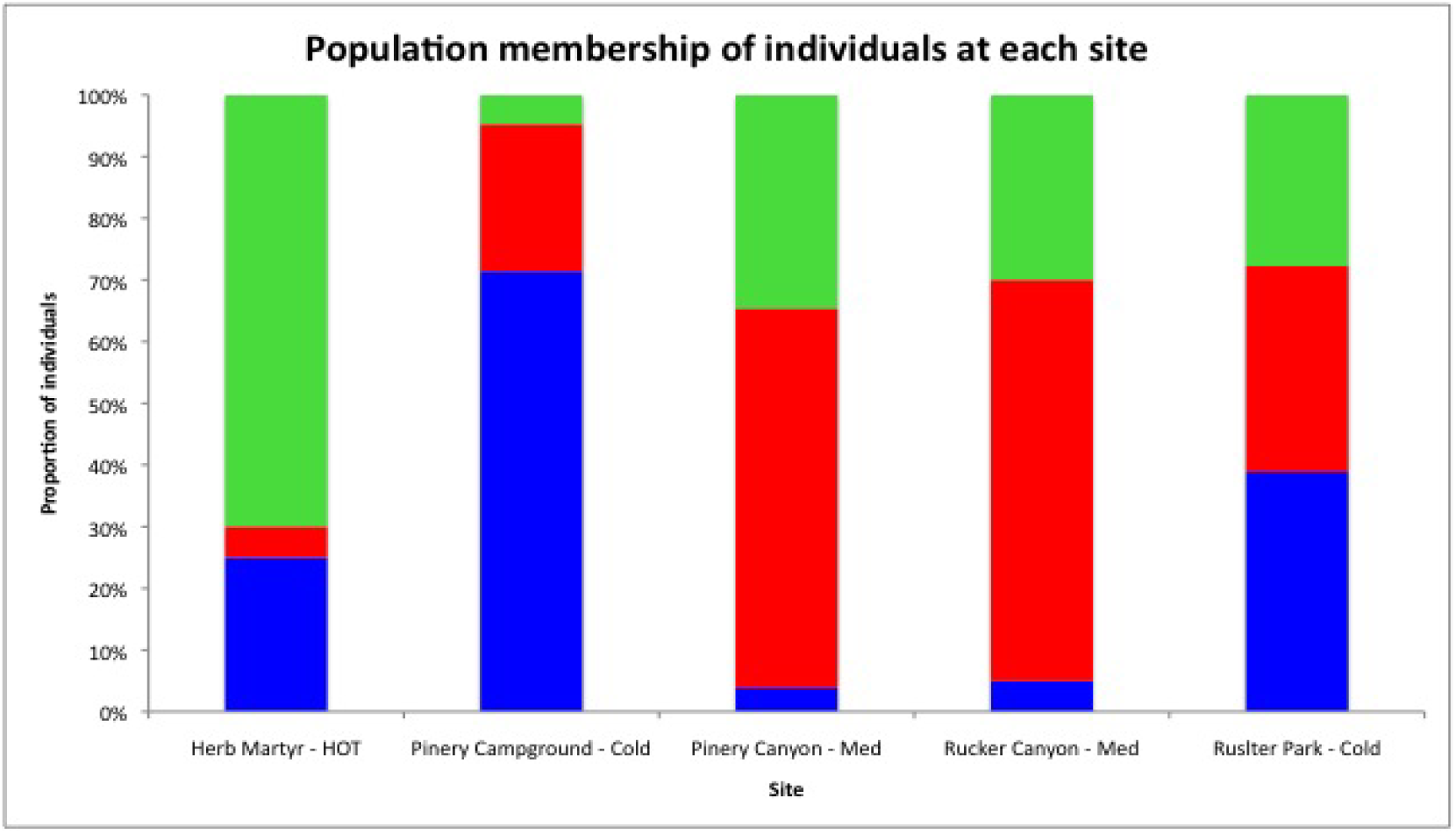
Population membership of individuals at each site. Population membership of individuals at each site. Herb Martyr is composed in majority of individuals from K1 genetic background (green). Pinery campground is composed in majority of individuals from K2 genetic background (blue), as is Ruslter park but to a lower extant. Pinery Canyon and Rucker Canyon are composed in majority of individuals from K3 genetic background (red).

### Isolation-by-temperature modulates gene flow patterns

To test whether geography, temperature, or dispersal phenotype (winged / wingless queen) influences gene flow patterns and genetic divergence within the Chiricahuas, we performed a series of analyses testing the interrelations between genetic divergence (Fst) and each of these factors. We first tested the simplest model of isolation-by-distance (IBD), where gene flow is only limited by geography. We performed a Mantel test between the Fst matrix within the Chiricahuas and geographic distance, and we did not find an effect of geographic distance on genetic divergence (Table 4). This indicates that within a Sky Island, gene flow is restricted, or promoted, by factors other than geographic distance between sites. Second, given the possible adaptation of *Monomorium* populations to temperature and the results from STRUCTURE, we tested a model of isolation-by-environment (IBE), where gene flow is limited by the temperature gradient. We performed a partial Mantel test between the Fst matrix and temperature distance between sites, while controlling for differences in elevation. We found a strong (0.64) and highly significant (p=0.008) effect of temperature on genetic divergence (Table 4), indicating that gene flow between sites with temperature differences is restricted.

**Table 4:** Mantel tests for IBD, IBE and IBA. We tested for IBD, IBE and IBA using mantel and partial mantel tests. We found a significant association between genetic differentiation and average annual temperature, suggesting that *M. emersoni* population structure is shaped primarily by differences in temperature across habitats, rather than absolute geographic distance, or potential for dispersal (queen phenotype).

| Variables tested (Variable controlled for) | Process tested | p-value |
| --- | --- | --- |
| Geographic distance vs Fst | IBD | 0.799 |
| Elevation vs Fst | IBD | 0.619 |
| Annual Temperature vs Fst (Elevation) | IBE | **0.008** |
| Dispersal potential vs Fst (Elevation) | IBA | 0.750 |
| Dispersal potential vs mtDNA distance (Geographic distance) | IBA (Mito) | 0.227 |
| Dispersal potential vs Annual Temperature (Elevation) | Queen vs Temp | 0.55 |

Third, we have also described a repeated adaptation of *M. emersoni* queen dispersal phenotype (winged or wingless) on each of the Arizona Sky Islands (*49*). We therefore tested a model of isolation-by-adaptation (IBA), where gene flow patterns are associated with dispersal potential between the sites. We define dispersal potential as the average winged queen frequency of the two sites being compared, which reflects how many winged queens are present to mediate dispersal between the two sites. We performed a Mantel test between Fst matrix and potential for dispersal between each pairwise comparison, while controlling for elevation, geographic distance, temperature, or latitude (Table 4). We did not find any significant isolation-by-adaptation in any of the tests (Table 4), indicating that the presence of winged queens does not influence the amount of gene flow occurring between two sites. Controlling for the temperature differences between sites did not improve the model, indicating that dispersal potential does not explain gene flow patterns within a Sky Island, even between sites of similar temperature. The absence of an association between a dispersal phenotype and gene flow patterns, even when controlling for covariates, is unexpected. It indicates that the achieved dispersal of the individual or its offspring is dissociated from the actual dispersal phenotype of the individual. Together, these result suggest that gene flow within a Sky Island is mostly shaped by the local adaptation to temperature rather than the dispersal phenotype of the queens.

## Discussion

Using an ecological genomics approach, we reveal that regions of *M. emersoni* genome show divergence between populations in a pattern consistent with natural selection caused by temperature. We show that this adaptation appears to influence gene flow patterns within a Sky Island, and that surprisingly, gene flow is not associated with queen dispersal phenotype. These seemingly contradictory observations may be explained by several possible scenarios: (*1*) Winged queens are indeed present, but do not perform dispersal between sites. Given the very high cost of developing wings and wings muscles (*55*, *77*, *78*), this scenario is unlikely and would be highly costly for the population and probably rapidly lost. (*2*) Wing phenotype could be plastic to some extant, and influenced by an environmental factor other than temperature. A dissociation between the achieved gene flow between two sites (Fst) and the dispersal potential (percent of winged queens in the two sites) could be explained by winged queens performing dispersal (gene flow), but giving rise to offspring that are wingless when sensing the right environmental cue. Such an effect could change the observed queen phenotype frequencies without affecting gene flow patterns between sites. We indeed previously found an association between queen phenotype and habitat fragmentation in *M. emersoni* (*49*), indicating that the phenotype is likely adaptive to some ecological variable. In the light of these results and our previous findings, we propose that the determination of the *M. emersoni* queen phenotype is environmentally plastic and may be largely influenced by environmental cues on the Chiricahuas Sky Islands.

Caste polyphenism is a nearly universal feature of ants that highlights their potential for the evolution and maintenance of environmentally induced morphologies (*79*, *80*). Most ant species show wing polyphenism, meaning that the determination of winged queens or wingless workers is controlled through an environmentally-controlled switch-like mechanism that is mediated by juvenile hormone (*43*, *79*, *80*). Wing alternative phenotypes in reproductive castes of some ant species have been found to be under the control of environmental factors (*81*, *82*). In *M. emersoni*, using an eco-evo-devo approach, we previously showed that the queen phenotype is found in fragmented and high elevation habitats, and postulated that an ancestral developmental potential of *Monomorium* queens for wing loss has facilitated the repeated adaptive evolution of wingless queens in *M. emersoni* and its associated gene regulatory network (*49*). Here, using an ecological genomics approach, we show evidence for a repeated adaption of *M. emersoni* to temperature gradients in the Arizona Sky Islands following the Pleistocene glaciation, and that this adaptation in the ecologically heterogeneous landscape has shaped gene flow patterns within a Sky Island.

It has been postulated that gene flow between different selective regimes may be promoted by the presence of phenotypic plasticity that facilitates phenotypic adaptation to the diverging ecological conditions (*75*). Organisms living in ecologically heterogeneous landscapes disperse across divergent environments where some of their adaptive phenotypes are advantageous, while other phenotypes may not be a good match for the new environment. In such a case, phenotypic plasticity can facilitate reaching a new optimal phenotypic combination of traits that matches the newly colonized environment (*83*–*85*). Indeed, within a single population, the number ecologically divergent loci that can be maintained by natural selection is limited (*74*, *86*), and a mismatch in the combination of niche specific traits can reduce fitness of individuals (*87*). Phenotypic plasticity can therefore facilitate the coexistence of multiple optimal adaptive phenotypes within a population (*83*, *84*).

In *M. emersoni,* synthesizing data from both eco-evo-devo and ecological genomics, we raise the possibility that there exists a rugged fitness landscape, where wing polyphenism and temperature tolerance both play a role in the ecological adaptation of *M. emersoni* to climate change in the Arizona Sky Islands (Figure 6). For each individual queen, the optimal phenotypic state at each of these two traits depends on the local environment, where local selection pressures may change rapidly over short geographical scales. A good match between the heterogeneous environment and the two distinct adaptive phenotypes (wing and temperature) can be attained by the combination of a genetic adaptation to temperature, which influences gene flow between sites, and an immediate plastic modulation of the offsprings dispersal phenotype to match more adequately its dispersal abilities to the environment where it now inhabits.

**Figure 6:**
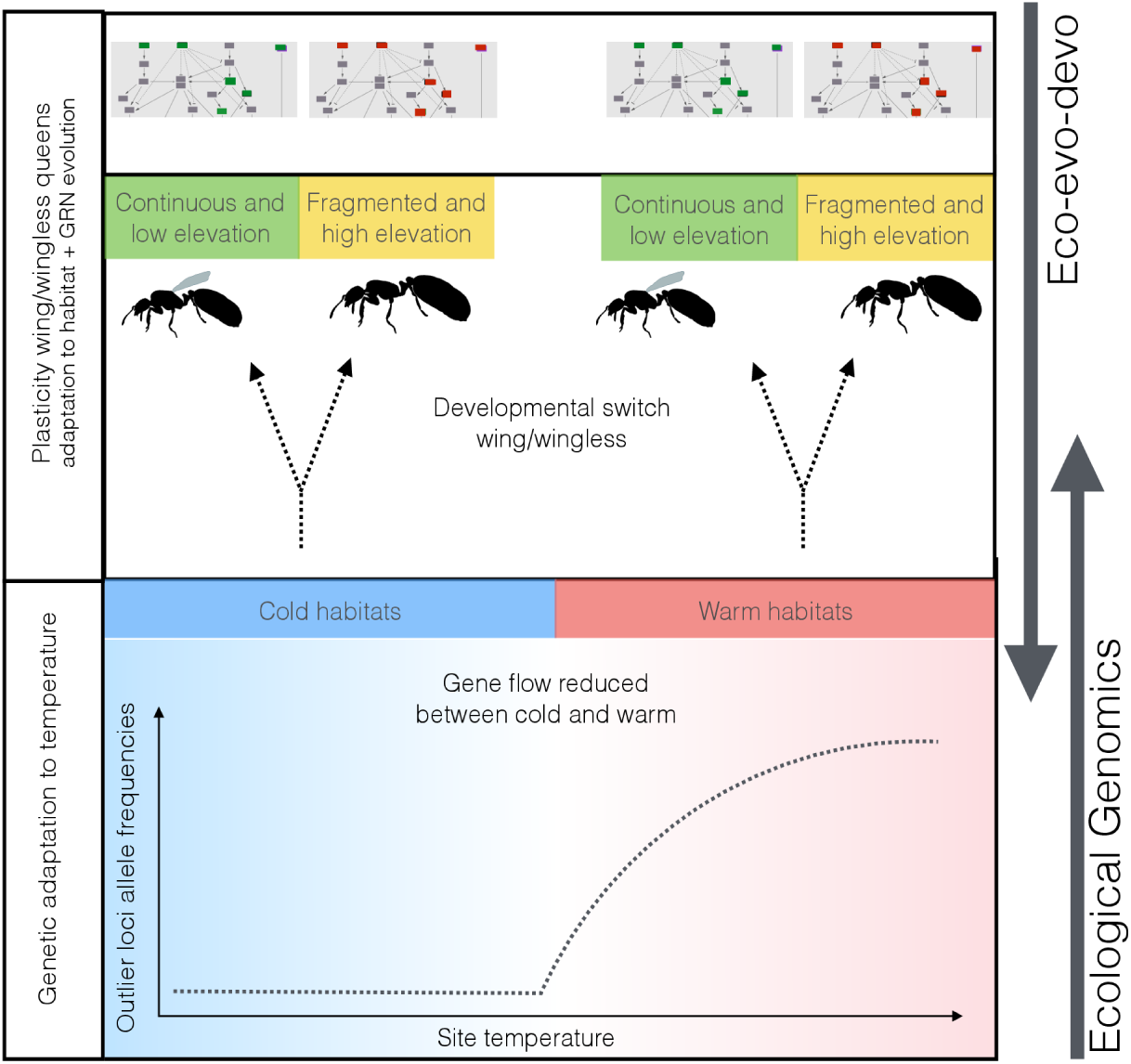
Adaptation in heterogenous environments. We revealed a potential genetic basis for temperature tolerance in *M. emersoni* to the replicated temperature gradients in the Arizona Sky Islands. Populations at collection sites with higher temperature show increased allelic frequencies of outlier loci (Lower box, representing Fig. 3). This genetic divergence associated with temperature tolerance is repeated on each of the Sky Islands, suggesting recurrent independent adaptation. Temperature divergence between habitats modulates gene flow patterns of winged queens within a Sky Island. During development (dotted arrows), queens of *M. emersoni* will develop into a winged or a wingless queen (Ant outlines). Because of the absence of a relationship between gene flow and dispersal potential, the association of wing phenotype and habitat fragmentation, and the ancestral developmental potential to produce alternative queen phenotypes, we propose that the queen wing phenotype in *M. emersoni* is likely under environmental influence (plastic). The gene regulatory network (Top box, and Fig. 2C) underlying wing development is interrupted in wingless queens (*49*), and shows similar interruption patterns within a Sky Island. Gene regulatory network are represented for each queen phenotypes in the top box. Genes highlited in green show conserved expression, and genes in red show interrupted expression in wingless queens (expression pattern that is different from the winged queens and *Drosophila melanogaster*. Within a Sky Island, the interruptions of the wing regulatory gene network in wingless queens are similar across sampling sites, but interruptions show differences between different Sky Islands (*49*).

### Evolutionary strategies following environmental change in replicated and heterogeneous landscapes

Environmental change imposes challenges to organisms during which they must either disperse, adapt, or adjust their phenotypes to the new environments (*6*, *88*, *89*). Synthesizing data from eco-evo-devo and ecological genomics, we show that *M. emersoni* possibly adopted both strategies repeatedly to survive climatic change in the Arizona Sky Islands; after the last glaciation, the once continuous population of *M. emersoni* was divided into isolated populations on each of the Sky Islands. *M. emersoni* moved to refugias in the cooler, higher elevation habitats that resemble most the habitat that was found at lower elevations during the glaciation (*50*) and that they were adapted to. In addition, *M. emersoni* could persist in the warmer lower mid-elevations of the Sky Islands by adapting to the warmer temperature. This can now be observed by the allelic frequencies of loci in islands of genomic divergence matching the temperature gradient, heat-shock tolerance of individuals coming from warmer habitats, and restricted gene flow between habitats divergent for temperature. Furthermore, this temperature adaptation occurred repeatedly across replicated populations: *M. emersoni* from each of the 5 Sky Islands we studied show outlier loci frequencies that are associated with temperature, and heat-shock tolerance was observed in colonies coming from different Sky Islands (*49*). The same loci across populations are repeatedly associated to the temperature gradient, suggesting a repeated and predictable evolution of temperature adaptation across independent replicates. However, a more detailed genomic mapping of the loci responsible for this adaptation may reveal more subtle differences across replicates that we were not able to identify with our analyses.

Simultaneously, *M. emersoni* queens also adjusted to the high elevation and fragmented habitats post-glaciation by repeatedly activating an ancestral developmental potential for wing loss in queens (*44*, *49*). We found that the underlying wing patterning gene network repeatedly evolved similar changes across Sky Islands in wingless queens, but also shows differences in expression across Sky Islands that are unpredictable and that are contingent on population history. Here, we suggest that the wing phenotype in queens may be influenced by the environment, and that such phenotypic plasticity may facilitate gene flow within each Sky Island, across different habitats. We raise the possibility that the maintenance of genetic temperature adaptation of *M. emersoni* in the Sky Islands was in part facilitated by the presence of an environmentally determined queen phenotype, which relaxed genetic constraints on adaptation across ecologically divergent environments. A relaxation of the genetic constraints imposed by ecologically divergent adaptations could explain the coexistence of a mosaic of different phenotypes within the same individual that best matches its environment (*75*, *85*). On rugged fitness landscapes, phenotypic plasticity may help jump across valleys of low fitness by immediately matching the phenotype to the environment an individual disperses to (or its offspring) (*85*), and therefore optimizing multiple adaptive traits simultaneously.

### Where is the locus of adaptation?

Experimental studies seeking to identify the locus of adaptation have been numerous (*8*,*12*,*14*,*19*,*20*,*23*,*43*, *90*–*92*), and an emerging question concerns our ability to discover adaptive genetic loci given the particular genetic architecture of each trait of interest (*35*, *93*). Past demographic events, population stratification, genetic pleiotropy, recombination rate variability, and phenotypic plasticty can alter our ability to identify loci underlying adaptations (*36*, *94*). More specifically, phenotypic plasticity and gene-by-environment interactions are likely to be pervasive in nature, and they can modulate the associations between the genotype and the phenotype (*35*, *36*, *94*). Our quest of revealing the genetic mechanisms and architecture underlying adaptive phenotypes will likely benefit from a combination of top-down and bottom-up approaches by leveraging their respective strengths. Organisms face a multitude of novel ecological pressures during climatic change, from changes in physical factors (temperature, precipitation), changes in species interactions, and changes in ecological communities (88,89,95,96). The selective environment often presents more than one ecological challenge at a time, and there may exist a variety of adaptive strategies adopted by a population living in a heterogeneous environment to persist in the face of environmental change. In the last decades, considerable attention has been given to demographic and genetic mechanisms to understand how species cope with climatic changes, but the role of development has been largely ignored. Integrating results from both approaches may reveal evolutionary strategies adopted by populations that would not be discovered by relying on only either of those approaches alone.

## Acknowledgements

We thank Robert A. Johnson for his indispensable help during field work and for discussions about the Arizona Sky Islands and its ant fauna. for We thank N. Aubin-Horth, L. Nilson, A. Hendry, J. Marcus, D. Schoen, and the Abouheif lab members for providing comments on this manuscript, as well as for instigating discussions on ecological genomics and evo-devo perspectives on the genetic basis of adaptation. This work was supported by grants from Fonds de la Recherche du Quebec - Nature et Technologies, Canada Research Chairs program, and National Sciences and Engineering Research Council (NSERC) to E.A., NSERC postgraduate scholarship to M-J.F.

## Data Availability

The environmental data is available through the World Clim database. Files with genotyping and phenotypic data are available from authors upon request.

